# Repetition suppression for mirror images of objects and not Braille letters in the ventral visual stream of congenitally blind individuals

**DOI:** 10.1101/2024.09.23.614517

**Authors:** Maksymilian Korczyk, Katarzyna Rączy, Marcin Szwed

## Abstract

Mirror-invariance effect describes the cognitive tendency to perceive mirror-image objects as identical. Mirrored letters, however, are distinct orthographic units and must be identified as different. Mirror-invariance must be ‘broken’ to enable efficient reading. Consistent with this phenomenon, a small, localized region in the ventral visual stream, the Visual Word Form Area (VWFA), exhibits repetition suppression to identical and mirror pairs of objects but only to identical pairs of letters. The ability of congenitally blind individuals to ‘break’ mirror invariance for pairs of mirrored Braille letters has been demonstrated behaviorally. However, its neural underpinnings have not yet been investigated. Here, in an fMRI repetition suppression paradigm, congenially blind individuals (both sexes) recognized pairs of everyday objects and Braille letters in identical (’p’ & ’p’), mirror (’p’ & ’q’), and different (’p’ & ’z’) orientations. We found repetition suppression for identical and mirror pairs of everyday objects in the parietal and ventral-lateral occipital cortex, indicating that mirror-invariant object recognition engages the ventral visual stream in tactile modality as well. However, repetition suppression for identical but not mirrored pairs of Braille letters was found in the left parietal cortex and the lateral occipital cortex but not in the VWFA. These results suggest notable differences in reading-related orthographic processes between sighted and blind individuals, with the LOC region in the latter being a potential hub for letter-shape processing.

**Significance Statement:** Mirror invariance is a tendency to recognize rotated objects as identical. Letters are unique shapes as people learn to recognize mirrored letters (e.g., ‘b’ and ‘d’) as distinct objects. In our study, we investigated the neural underpinnings of tactile mirror invariance in congenitally blind individuals. We demonstrated engagement of the parietal, occipital, and ventral visual regions in mirror-invariant tactile object recognition, indicating that this perceptual bias extends beyond the visual modality. Moreover, we found that unlike in the sighted, it was the parietal and lateral occipital cortex that showed neural signatures of breaking mirror invariance for Braille letters in congenitally blind individuals, suggesting substantial differences between visual and tactile reading.

## 1 Introduction

Mirror invariance is an automatic predisposition of the visual system to recognize objects reflected on both vertical and horizontal axes as identical (Corballis & Beale, 1976; Tarr & Pinker, 1989). The neural underpinnings of this phenomenon were demonstrated in both macaque monkeys (Rollenhagen & Olson, 2000) and humans (e.g., Dehaene et al., 2010; Pegado et al., 2011) using fMRI repetition suppression studies. Repetition suppression – known as well as an fMRI priming effect - refers to a decrease in neural response to a target stimulus (e.g., the word “SHEEP” or a picture of a sheep) when it is preceded by an identical stimulus (referred to as a prime). Mirror-invariance in object recognition is reflected in repetition suppression of the neural response to a target object in regions of the ventral visual stream when it is preceded by its identical or mirrored image, even though the retinal projections for this object and its mirror reflection differ significantly (Dehaene et al., 2010; Pegado et al., 2011; Dilks et al., 2011). Mirror-invariance, however, poses a significant challenge when children start to read, as mirrored letters of many scripts, such as the Latin ’b’ and ’d,’ have to be discriminated as different (Cornell, 1985; Fischer & Koch, 2016; Ahr et al., 2016). Once mirror invariance for letters is ’broken,’ the visual word form area (VWFA) – a region crucial for reading (Zhan et al., 2023) – shows repetition suppression for mirrored pairs of objects but not for mirrored letters (Dehaene et al., 2010; Pegado et al., 2011).

The Braille alphabet consists of mirror letters as well. In fact, it has even more of them than the Latin alphabet. Blind children, just like sighted children, struggle with mirror letters (Millar, 2004) when they start to learn tactile Braille. De Heering and colleagues (2018) demonstrated that analogously to the sighted, congenitally blind individuals also ’break’ mirror invariance for Braille letters, since analogously to the sighted readers, judging the shape of two mirrored letters as the same is more demanding for them than doing so for objects. In another behavioral study, we (Korczyk et al., 2024) observed, similarly, high expertise of blind individuals with Braille letters, that is, their proficiency in recognizing different orientations of mirror letters. At the same time, however, we did not observe any significant difficulties when blind participants discriminated against left-right oriented Braille letters as having the same shape. We speculated that blind readers of Braille might exhibit greater selective attention and thus filter task-irrelevant information more efficiently (Korczyk et al., 2024). Nevertheless, both studies suggest considerable perceptual expertise in processing Braille letters in blind individuals.

Here, we investigated the neural underpinnings of mirror-letter discrimination in Braille reading with an fMRI adaptation paradigm. Participants were presented with two categories of haptic stimuli: everyday objects and Braille letters (Fig 1A). All stimulus categories were presented sequentially in three different prime-target pairs: identical (’b’ and ’b’), mirror (’b’ and ’d’), or different (’b’ and ’g’) (Fig. 1B). We reasoned that if we find identity and mirror-priming for objects and identical but no mirror-priming for Braille letters in a given brain region, it will provide evidence that this brain region is the locus of letter shape processing in blind individuals. If that brain region is found in the location of the sighted VWFA, it would support the VWFA being a task-specific reading region of the brain (Amedi et al., 2017), independent of the input modality, and not a general language area (Bedny, 2017). However, if we find this pattern of results in, e.g., the lateral occipital cortex (LOC), which has been suggested to process shape information independent of the input modality (Hannagan et al., 2015), it would suggest that blind individuals’ reading system notably differs from the one in the sighted.

**Figure 1.**
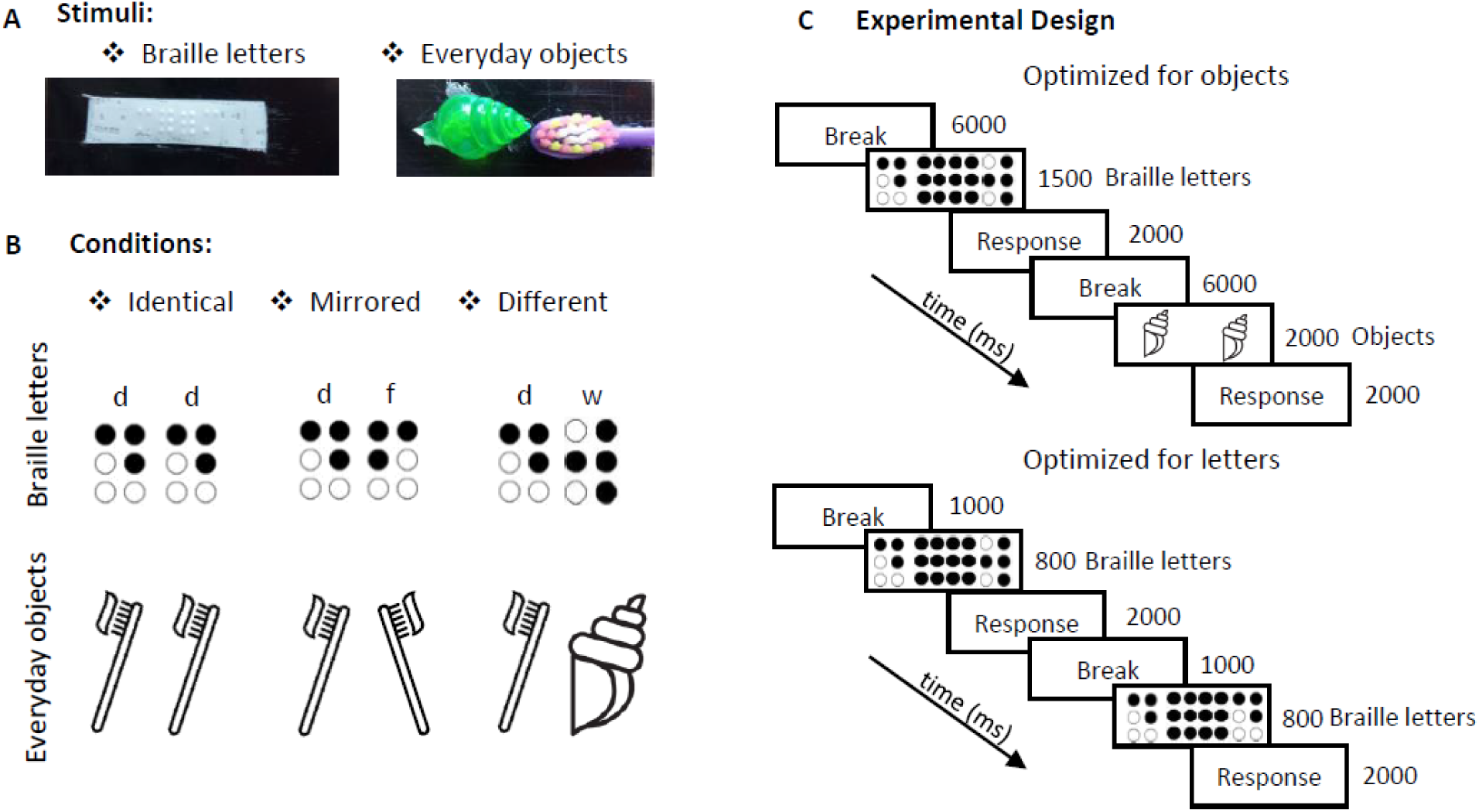
**(A)** Tactile stimuli used in the first fMRI experiment: Braille letters and everyday objects. Both Braille letters and everyday objects were presented on a conveyor belt. **(B)** In both fMRI experiments, all stimuli were presented under identical, mirrored, and different conditions. **(C)** Experimental design. An fMRI priming paradigm was implemented. In the first fMRI experiment, Braille letters were presented for 1500 ms and objects for 2000 ms. This presentation timing was optimal for objects but not for Braille letters. In the second fMRI experiment, the focus was on Braille letters, presented for 800 ms, which was optimal for Braille letter presentation.

## 2 Materials and Methods

In the current study, we conducted two fMRI experiments. The first was optimized for perception of 3D everyday objects, with longer stimulus presentation times (referred to as Experiment 1). The second was optimized for perception of Braille letters, with shorter stimulus presentation times (referred to as Experiment 2) (Fig. 1C).

### 2.1 Participants

Eighteen congenitally blind native Polish speakers (7 females, mean age: 31.9 years, SD = 5.3) participated in the first fMRI experiment. In the second fMRI experiment, eighteen congenitally blind Polish speakers also took part (10 females, mean age: 33.2 years, SD=6.14). Fourteen of the eighteen subjects participated in both fMRI experiments. The congenitally blind individuals are a hard-to-find, rare clinical population, and such a sample size is comparable to, or larger than recent fMRI studies with blind individuals [e.g., 17 participants in Kanjlia, Feigenson, & Bedny (2021); 8 participants in Vetter and colleagues (2021); 9 participants in Bola et al. (2023)].

In the first fMRI study, the main causes of blindness among the participants included retinopathy of prematurity (n = 12), atrophy of the optic nerve (n = 4), and other causes (n = 2) (for a detailed description, see Table 1). Eleven out of 18 participants were completely blind, while the others had primitive sensitivity to light. In the second fMRI study, the main causes of blindness among the participants included retinopathy of prematurity (n = 13), atrophy of the optic nerve (n = 3), and other causes (n = 2) (for a detailed description, see Table 2). Thirteen participants were completely blind, while the others had primitive sensitivity to light.

**Table 1.** Information about congenitally blind individuals. First fMRI experiment

| Participant | Age (Years) | Gender | Cause of Blindness | Reading Hand |
| --- | --- | --- | --- | --- |
| 1 | 45 | Female | Retinopathy of prematurity | right (index finger) |
| 2 | 32 | Female | Retinopathy of prematurity | right (index finger) |
| 3 | 32 | Female | Retinopathy of prematurity | left |
| 4 | 31 | Male | Retinopathy of prematurity | right |
| 5 | 35 | Male | Glaucoma | right |
| 6 | 24 | Male | Atrophy of the optic nerve | left (index finger) |
| 7 | 30 | Female | Atrophy of the optic nerve | right |
| 8 | 34 | Male | Genetic | left |
| 9 | 30 | Male | Atrophy of the optic nerve | right |
| 10 | 28 | Female | Retinopathy of prematurity | right |
| 11 | 27 | Male | Retinopathy of prematurity | right |
| 12 | 31 | Female | Atrophy of the optic nerve | right (index finger) |
| 13 | 25 | Male | Retinopathy of prematurity | left |
| 14 | 26 | Male | Retinopathy of prematurity | left |
| 15 | 36 | Male | Retinopathy of prematurity | left |
| 16 | 39 | Male | Retinopathy of prematurity | left |
| 17 | 39 | Female | Retinopathy of prematurity | left (index finger) |
| 18 | 31 | Male | Retinopathy of prematurity | left |

**Table 2.**
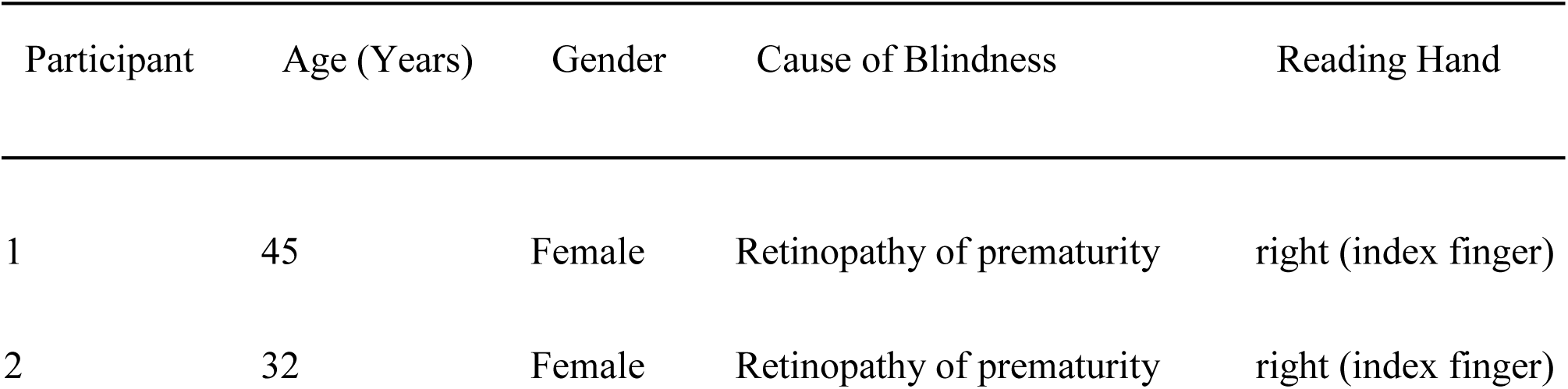
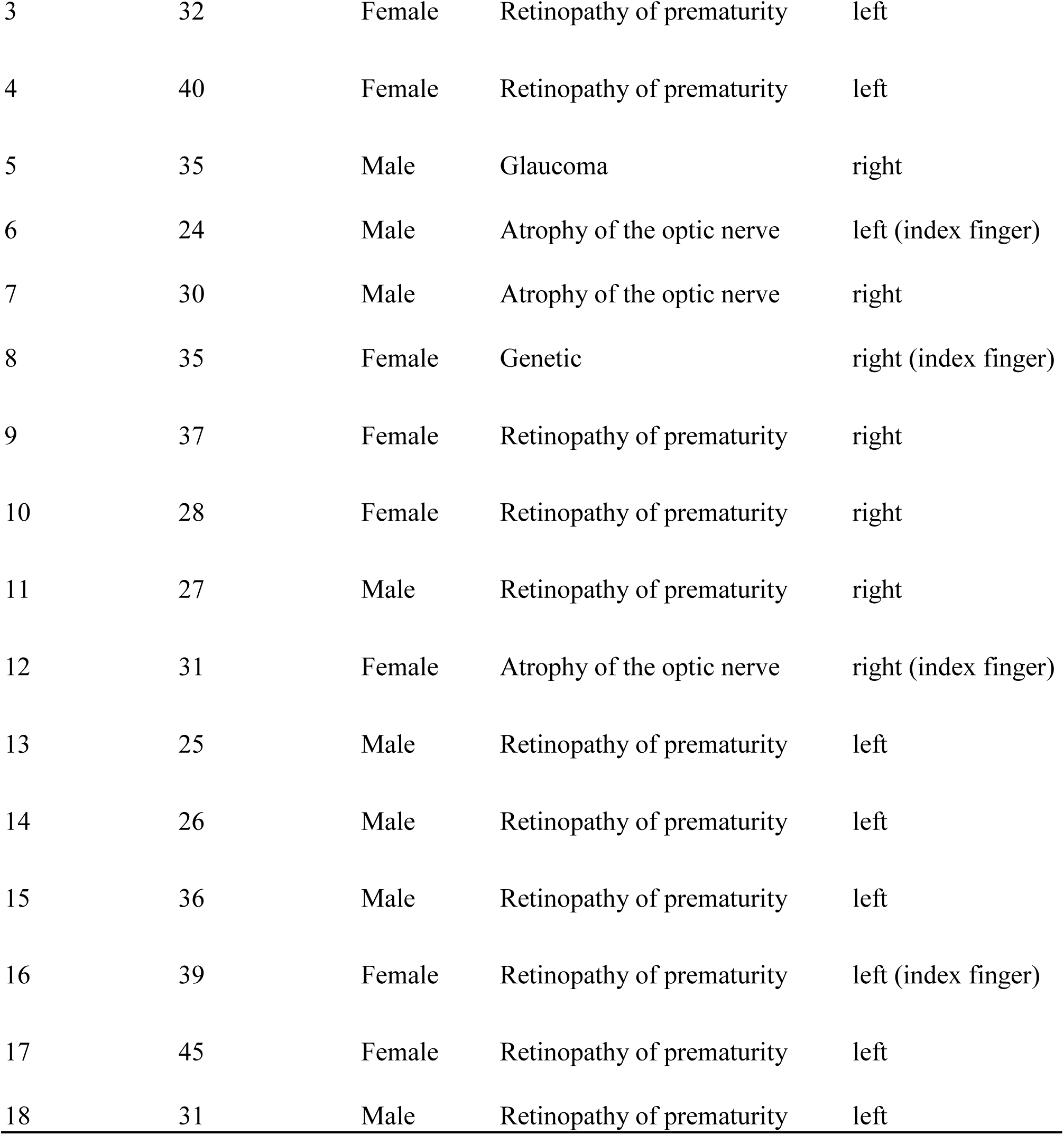
Information about congenitally blind individuals. Second fMRI Experiment.

| Participant | Age (Years) | Gender | Cause of Blindness | Reading Hand |
| --- | --- | --- | --- | --- |
| 1 | 45 | Female | Retinopathy of prematurity | right (index finger) |
| 2 | 32 | Female | Retinopathy of prematurity | right (index finger) |
| 3 | 32 | Female | Retinopathy of prematurity | left |
| 4 | 40 | Female | Retinopathy of prematurity | left |
| 5 | 35 | Male | Glaucoma | right |
| 6 | 24 | Male | Atrophy of the optic nerve | left (index finger) |
| 7 | 30 | Male | Atrophy of the optic nerve | right |
| 8 | 35 | Female | Genetic | right (index finger) |
| 9 | 37 | Female | Retinopathy of prematurity | right |
| 10 | 28 | Female | Retinopathy of prematurity | right |
| 11 | 27 | Male | Retinopathy of prematurity | right |
| 12 | 31 | Female | Atrophy of the optic nerve | right (index finger) |
| 13 | 25 | Male | Retinopathy of prematurity | left |
| 14 | 26 | Male | Retinopathy of prematurity | left |
| 15 | 36 | Male | Retinopathy of prematurity | left |
| 16 | 39 | Female | Retinopathy of prematurity | left (index finger) |
| 17 | 45 | Female | Retinopathy of prematurity | left |
| 18 | 31 | Male | Retinopathy of prematurity | left |

All participants were blind from birth and had never experienced any patterned vision. None of the participants reported any additional sensory, motor disabilities, neurological, or psychiatric problems.

The Jagiellonian University Ethics Committee approved both fMRI studies. All participants provided written informed consent before the study and were reimbursed for taking part in both experiments separately.

### 2.2 Stimuli

In the first fMRI experiment, two categories of stimuli were used: everyday objects (14: the lid of a tic-tac mint box, ½ adhesive tape, measuring spoon (7.5 mm), toothbrush, pen lid, bag clips, a measuring spoon (5 mm), three types of acrylic plastic seashells for aquariums, four pills in a blister pack, ear stick (q-tip), comb, and straw) and Braille letters (8: d, e, h, m, p, r, y, and z) (Fig. 1A). All stimuli were presented in pairs under three conditions: identical (identical pairs of stimuli presented in the same orientation), mirrored (identical pairs of stimuli presented in mirror symmetry), and different (different pairs of stimuli presented in different orientations) (Fig. 1B). All stimuli were glued to a plastic plate (8×3 cm). A one-centimeter distance between pairs of everyday objects was maintained. All everyday objects were purchased from common supermarkets, drugstores, or stationery shops. Before the fMRI experiment, we conducted pilot and behavioral experiments (Korczyk et al., 2024) to verify whether the subjects could accurately and quickly recognize the everyday objects later used in both behavioral and fMRI experiments. This allowed us to select fourteen stimuli that all participants correctly recognized in less than 2 seconds.

Braille letter stimuli were printed on an RL-350 Braille labeler on Reizen vinyl label tape. Between pairs of Braille letters, we added two six-dot letters, and these stimulus prints were then glued in the center of the 8×3 cm plastic plate.

In the second fMRI experiment, the primary focus was on Braille letters, and therefore, we presented the participants with all 32 letters of the Polish alphabet displayed in pairs in three conditions: identical, mirror, and different (see above).

### 2.3 Procedure

In the first fMRI experiment, the everyday objects and Braille letters were presented on a conveyor belt designed for tactile stimuli. We used Presentation software (Neurobehavioral Systems; https://www.neurobs.com/) to present auditory cues to control the presentation of the tactile stimuli. Each cue instructed the experimenter to move the conveyor belt to the next set of stimuli. The conveyor belt was custom-designed to be fMRI-compatible (Neurodevice, Warsaw, Poland) for this study (see Supplementary Fig. 1A). It featured 80 slots for attaching plastic plates with stimuli. A total of 39 plastic plates, used for all runs, were attached to every second slot. To prevent potential errors, we introduced spacers (gaps) between each stimulus plate (Supplementary Fig. 1B).

The device was positioned above the participants’ thighs (Supplementary Fig. 1C). The participants’ reading hand was placed in a designated aperture, allowing them to touch only one pair of stimuli at a time (see Supplementary Fig. 1D). Participants read Braille and recognized everyday objects using their preferred Braille reading hand. Throughout all runs, the movement of the conveyor belt was controlled by the same researcher.

The first fMRI experiment consisted of seven runs, and participants spent 90 minutes inside the scanner to complete all tasks. Across all runs, we presented 138 pairs of everyday objects and 135 pairs of Braille letters. In odd-numbered runs (1, 3, 5, and 7), we presented 18 pairs of Braille letter stimuli (6 pairs of each condition) and 21 pairs of everyday objects (7 pairs of each condition). In even-numbered runs (2, 4, and 6), we presented 21 pairs of Braille letter stimuli (7 pairs of each condition) and 18 pairs of everyday objects (6 pairs of each condition). Each trial began with auditory information indicating the type of stimuli to be presented, followed by an auditory cue (a 200-msec beep at a frequency of 4000 Hz), which signaled either a trial with Braille letters (presented for 1500 ms) or a trial with everyday objects (presented for 2000 ms). A second auditory cue (a 200-msec beep at a frequency of 44100 Hz) signaled the end of the trial. To minimize top-down processing, we employed a catch trial paradigm during which participants focused on recognizing either a seashell or the letter ’M.’ In Polish, the word for seashell (muszelka) starts with the letter ’M,’ making the task more accessible for the subjects. Participants had 2000 ms to indicate (by pressing a corresponding button) whether they recognized 1 seashell or 1 letter ’M,’ or 2 shells or 2 letters ’M,’ respectively (see Fig. 1C). Following this, there was a 6000 ms break for the researcher to move the belt in the conveyor belt device.

In the second fMRI experiment, stimuli were also presented using the Presentation software (Neurobehavioral Systems; https://www.neurobs.com/). The Braille letters were displayed on an fMRI-compatible Braille display (Neurodevice, Warsaw, Poland, Debowska et al., 2013), similar to commercial Braille devices, featuring pneumatically driven Braille pins. This device can simultaneously present five Braille letters, allowing participants to read them in a manner identical to reading regular Braille text. The Braille display was positioned on the participants’ thighs, on the side corresponding to their reading hand.

The second fMRI experiment consisted of five functional runs. In each run, we presented 108 pairs of Braille letters (36 pairs of each condition) along with 16 ’blank trials’, during which no auditory cue or tactile stimuli was presented on the Braille display for 2800 ms (the same duration as the trial). Each trial began with an auditory cue (a 200-msec beep at a frequency of 1500 Hz), indicating a trial with Braille letters, which were presented for 800 ms. Each pair of Braille letters was separated by a double six-dot character (in the Polish Braille alphabet, this character has no meaning). Subsequently, participants heard another auditory cue (a 200-msec beep at a frequency of 1500 Hz), signaling the end of the trial. During this experiment, to minimize top-down effects, participants were instructed to press one button when they recognized a pair of three-dot Braille letters (e.g., “l”), another when they perceived only one three-dot Braille letter in a pair, and to refrain from pressing any button when no three-dot Braille letters were presented. They had 2000 ms to respond (see Fig. 1C), followed by a 1000 ms break.

### 2.4 fMRI Data Acquisition

All fMRI data were acquired at the Małopolskie Centrum Biotechnologii in Kraków. Both fMRI experiments followed the same settings for functional and anatomical scans. Functional MR scans were collected using an EPI sequence on a 3T Siemens Skyra scanner equipped with a 64-channel head coil (flip angle = 70°; TR = 1200 ms; TE = 27 ms; FOV = 240 mm; matrix size = 80 x 80). Forty-two contiguous axial slices with a thickness of 3.0 mm and an in-plane resolution of 3.0 x 3.0 mm² were acquired. For anatomical reference and spatial normalization, T1-weighted images were acquired using an MPRAGE sequence (208 slices; FOV = 250 mm; TR = 1800 ms; TE = 2.37 ms; voxel size = 0.9 x 0.9 x 0.9 mm³).

### 2.5 Behavioral data analysis

In both experiments, we implemented a catch-trial paradigm, and thus, accuracy was measured. Reaction time, however, was not assessed as neither the priming effect nor the effect of stimulus orientation would be observable in any of the tasks. The participants’ ability to adequately recognize stimuli and their orientation was assessed during our behavioral study (Korczyk et al., 2024) with the same participants.

### 2.6 fMRI data analysis

All fMRI data were analyzed using the SPM12 software package (https://www.fil.ion.ucl.ac.uk/spm/software/spm12/). Data preprocessing steps included: (1) slice timing correction; (2) realignment of all EPI images to the first image; (3) coregistration of the anatomical image to the mean EPI image; (4) normalization of all images to MNI space; and (5) spatial smoothing with a 6-mm FWHM kernel. The hemodynamic response function (HRF) for main predictors (identical, mirror, and different pairs) and six estimated movement parameters as confound predictors were first modeled within a general linear model (GLM, Friston et al., 1997) for each participant.

#### 2.6.1 Whole Brain Analysis

Next, for both everyday objects and Braille letters, we conducted the following comparisons: (1) mirror condition versus identical condition, (2) different versus identical condition, and (3) different versus mirror condition. Subsequently, we overlapped brain activation induced by two different comparisons: (1) different versus identical and different versus mirror conditions for everyday objects; and (2) different versus identical and mirror versus identical conditions for Braille letters; (3) different versus identical conditions in both everyday objects and Braille letters.

Finally, we conducted a random-effects ANOVA analysis for the group. In the first experiment, we applied a voxel-wise threshold of p < 0.05 FWE and a cluster extent threshold of p < 0.05 FWE. In the second experiment, we used a voxel-wise threshold of p < 0.001 (uncorrected) and a cluster extent threshold of p < 0.05 FWE. To support the localization of the observed effects, we used a probabilistic atlas of the human brain implemented in the SPM Anatomy Toolbox 2.2b (Eickhoff et al., 2005).

#### 2.6.2 ROI analysis and functional localizer

To localize the regions involved in reading, we added an additional localizer run in our second fMRI experiment. We contrasted the activation induced by three Braille letters, each separated by six-dot letters, to five six-dot letters, which served as the basic tactile stimulus. This contrast revealed activation in the parietal-dorsal-ventral stream. Based on these results, we created anatomical masks located within this network.

The MarsBaR 0.44 Toolbox (Brett et al., 2002) (http://marsbar.sourceforge.net/) was used to conduct functionally guided ROI (Region of Interest) analyses. We examined signal changes induced by three different conditions (identical, mirror, and different) for everyday objects and Braille letters in both fMRI experiments. Anatomical masks for the left occipital-temporal and bilateral parietal areas were created using the SPM Anatomy Toolbox 2.2b (Eickhoff et al., 2005). The left occipital-temporal regions encompassed areas such as: 1) the primary visual cortex (V1 and V2, i.e., BA 17 and BA 18), 2) middle-temporal cortex (hOC5 (V5 / MT+)), 3) ventral (V3v / V4) and 4) dorsal extrastriate cortex (hOC3d / hOC4d), 5) fusiform gyrus (Areas FG1, FG2, FG3, and FG4), and 6) lateral occipital cortex (extrastriate areas hOc4la and hOc4lp) (Supplementary Fig. 2A). For the bilateral parietal cortex, we created masks for 1) the intraparietal sulcus (Areas hIP1, hIP2, hIP3), 2) motor cortex (Areas 4a and 4p), and 3) primary somatosensory cortex (Areas 1, 2, 3a, 3b) (Supplementary Fig. 2B).

These selections were based on relevant literature concerning visual mirror invariance and tactile symmetry in sighted individuals (e.g., Aspell et al., 2010; Dehaene et al., 2010; Pegado et al., 2011; Kitada et al., 2014; Fujimoto et al., 2017) and activations obtained in the localizer. Next, we extracted the beta estimates from each voxel within the anatomical masks and averaged them across voxels within the masks for each subject. We then entered the mean beta estimates into a repeated-measures ANOVA to assess the effect of orientation on the activation of these regions. All tests were corrected for multiple comparisons using the Bonferroni correction.

## 3 Results

In the first fMRI study, participants achieved an accuracy of 73.5% (SD = 13.7%), while in the second study, they achieved an accuracy of 76.6% (SD = 15.0%). The paired t-test showed no difference in performance obtained in both experiments t(17)=-0.617, p=0.545.

### 3.1 Experiment 1 - Everyday objects and Braille letters Whole-brain analysis

To probe for repetition suppression, we first contrasted everyday objects presented in different versus identical conditions (identity priming). We observed a repetition suppression effect bilaterally in a fronto-parieto-temporo-occipital network (Fig. 2). Specifically, the activation was observed bilaterally in the dorsal parietal regions, including the left inferior parietal lobule gyrus (−39, -40, 50; t=6.98, 367 voxels) and the right precuneus gyrus (45, 12, -70; t=6.26, 45 voxels), as well as the superior parietal lobule gyrus (24, -52, 56; t=5.85, 60 voxels), in the frontal regions, encompassing the left superior frontal gyrus (−24, -7, 53; t=5.64, 31 voxels) and the right posterior-medial frontal gyrus (6, 11, 53; t=5.79, 135 voxels), as well as in the occipitotemporal cortex, including the ventral visual stream with activations in the left fusiform gyrus (−33, -55, -10; t=7.34, 75 voxels) and the right fusiform gyrus (27, -70, -7; t=5.44, 14 voxels) (Fig. 2A and Supplementary Table 1).

**Figure 2.**
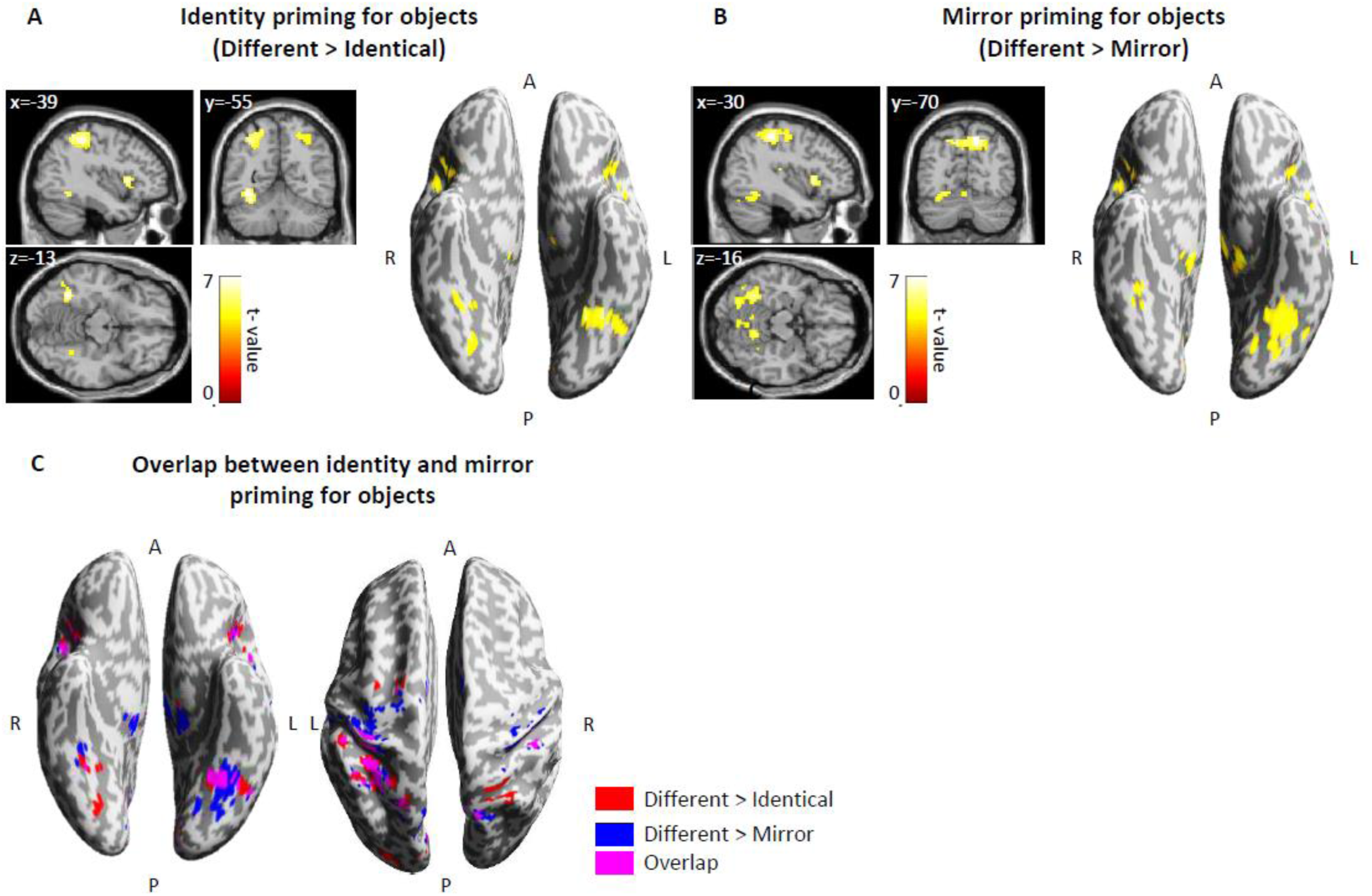
fMRI priming effects for tactile objects. **(A)** Identity priming: Objects induced bilateral effects in the dorsal parietal, posterior-medial frontal, and occipito-temporal regions. **(B)** Mirror priming: Objects evoked mirror priming in a wide area of the brain bilaterally, including the visual cortex, dorsal parietal, and posterior-medial frontal regions. **(C)** Overlap between identity and mirror priming for objects: We observed overlap in the left fusiform cortex (−33, -55, -19) and parietal cortex, including the intraparietal sulcus, motor cortex, and primary somatosensory cortex. However, mirror priming induced broader activation in this region compared to identity priming. Thresholds: **(A-C)** p < 0.05 FWE voxel-wise, p < 0.05 FWE cluster-wise. R-right, L-left, A-anterior, P-posterior.

Subsequently, we compared brain activation induced by everyday objects presented in the mirror versus identical conditions (mirror priming) and in the opposite direction (identical versus mirror conditions). We did not find any significant effects for either comparison, even using an exploratory cluster-wise threshold of p < 0.01, with a voxel-wise threshold of p = 0.01 (uncorrected).

Finally, we compared everyday objects presented in different versus mirror conditions (mirror priming). Similar to identity priming, we observed activations in a large portion of the bilateral fronto-parieto-temporo-occipital network. Specifically, activations were found bilaterally in the parietal cortex, including the left postcentral gyrus (−45, -28, 50; t=7.47, 586 voxels) and the right postcentral gyrus (48, -28, 53; t=6.38, 134 voxels), in the frontal regions, spanning the left posterior-medial frontal (−3, -7, 56; t=6.82) and the right posterior-medial frontal (6, 5, 53; t=6.95), both in one cluster with 257 voxels. Furthermore, activation was observed in the occipitotemporal cortex, including the ventral visual stream, with activations in the left fusiform gyrus (−33, -55, -13; t=6.45, 201 voxels) and the right fusiform gyrus (33, -43, -19; t=4.19, 115 voxels) (Fig. 2B and Supplementary Table 2). Activations for identity and mirror priming overlapped in the frontal and parietal cortex, as well as in the left fusiform cortex (with a sub-peak at coordinates -33, -55, -19) (Fig. 2C).

We repeated the analysis for Braille letters but did not observe any significant effects for either identity priming (different > identical) or mirror priming (mirror > identical). We speculated that the presentation times for Braille letters were too long to observe the priming effect, which is known to depend on stimulus presentation time (Schacter et al., 2004; Rączy et al., 2019). Hence, we performed a second fMRI experiment, optimizing the stimulus presentation time for Braille letters.

#### ROI analysis

First, we conducted repeated-measures ANOVAs (with 3 conditions: identical, mirror, and different) for everyday objects with beta estimates averaged across voxels within the left occipital-temporal regions as the dependent variable. This analysis was performed in various ROIs (including anatomical location of regions in the parietal, dorsal and ventral visual network). In all anatomical regions of the left occipital-temporal cortex, we obtained a main effect of orientation (all F’s > 9.78; all p < 0.001; η² between 0.36 and 0.59). In each anatomical ROI, pairs of everyday objects presented in the different condition induced significantly greater activation than those in the identical condition (all p < 0.037) or mirror conditions (all p < 0.003). There was no significant difference between the identical and mirror conditions (p > 0.377). Similar results were obtained when combining all regions into one ROI [F(2,34) = 18.78, p < 0.001; η2 = 0.53; post-hoc: different > identical p < 0.001; different > mirror p < 0.001, identical > mirror p= 1.000) (Fig. 4A).

**Figure 4.**
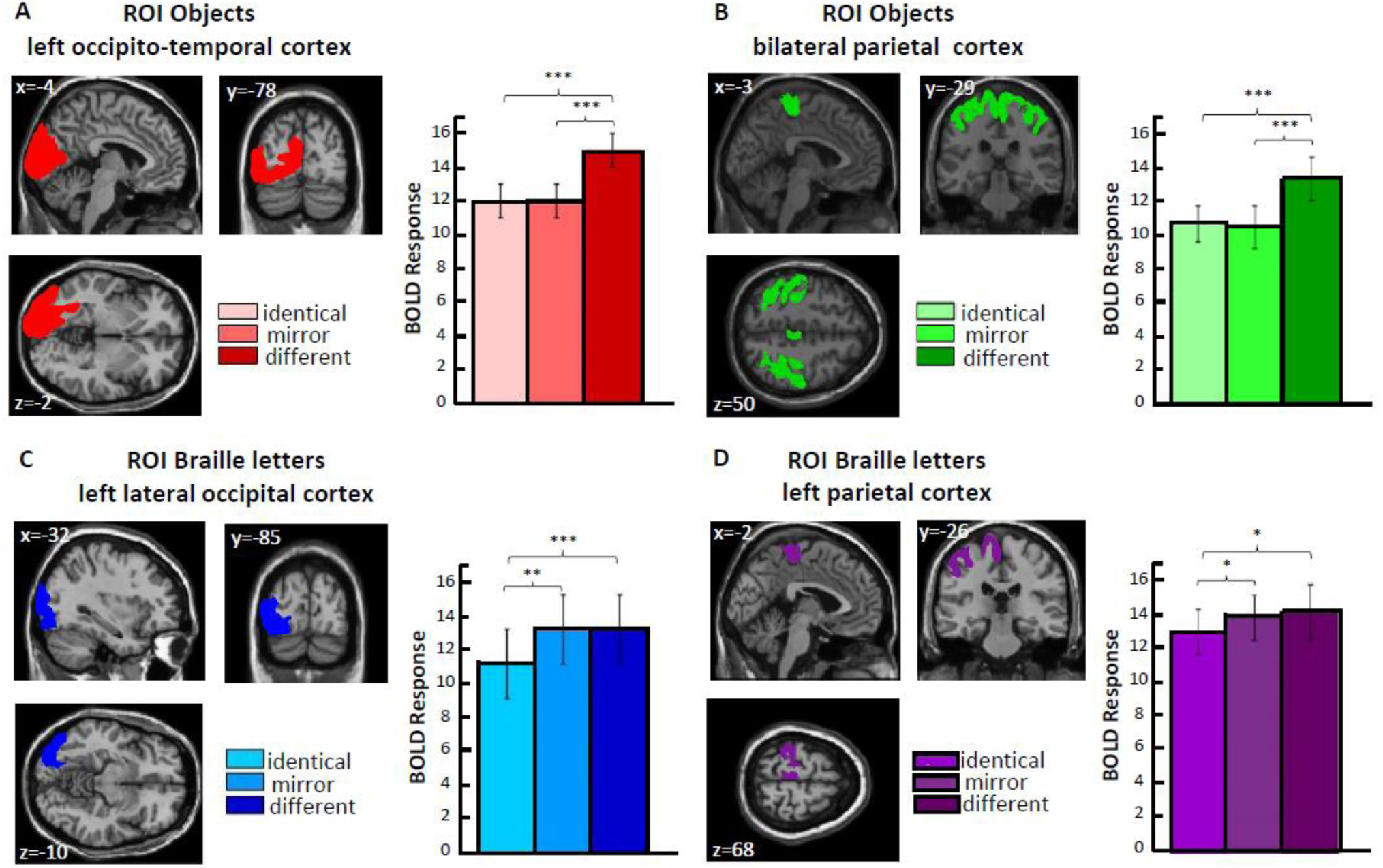
ROI results. **(A)** The analysis was performed in six anatomical ROIs within the left occipito-temporal regions for the signal change induced by three conditions (identical, mirror, and different) for objects in the first fMRI experiment. ROIs included: primary visual cortex (V1 and V2), middle-temporal cortex (V5), ventral and dorsal extrastriate cortex, fusiform gyrus, and lateral occipital cortex. In all ROIs, pairs of different objects induced significantly greater activation than pairs of same and mirror objects. **(B)** The same activation patterns were observed in all ROIs in the bilateral parietal cortex. ROIs included: the intraparietal sulcus, motor cortex, and primary somatosensory cortex. Both results **(A)** and **(B)** suggest that mirror invariance for everyday objects in cognitively blind individuals engages the brain in occipito-temporo-parietal regions. **(C)** We observed mirror discrimination for single Braille letters in the left lateral occipital cortex but not in the left ventral occipito-temporal cortex. **(D)** We observed mirror discrimination for single Braille letters in the left parietal cortex. Thresholds levels: *p < 0.05, ** p < 0.01, ***p < 0.001. Error bars represent S.E.M.

Next, the ROI analysis was repeated for the parietal regions. Repeated-measures ANOVAs (with 3 conditions: identical, mirror, and different) were performed on participants’ beta estimates averaged across voxels within the separate anatomical masks of parietal regions. The effect of orientation was significant in each ROI (all F’s > 14.62; all p < 0.001; η² between 0.46 and 0.66). In each ROI, pairs of everyday objects presented in the different condition induced significantly greater activation than those in the identical condition (all p < 0.001) or mirror conditions (all p < 0.008). No significant difference was found between the identical and mirror conditions (p = 1.00). Similar results were found when combining all regions into one ROI [F(2,34) = 28.06, p < 0.001; η2 = 0.62; post-hoc: different > identical p < 0.001; different > mirror p < 0.001, mirror > identical, p =1.000) (Fig. 4B).

### 3.2 Experiment 2 - Braille letters

In this second experiment, the presentation times were optimized for Braille letters, and only Braille letters were presented to the participants.

#### Whole-brain analysis

First, we contrasted Braille letters presented in different > identical conditions (identity priming). This contrast showed activations in the parietal cortex bilaterally [left: inferior parietal lobule (−42, -40, 4; t=6.00, 372 voxels); right: inferior parietal lobule (45, -37, 47; t=5.49, 158 voxels)], precentral gyrus (−42, 5, 29; t=3.86, 128 voxels), and in the left inferior occipital gyrus (−42, -64, -7; t=4.53, 70 voxels), as well as bilaterally in the occipital pole [left: -30, -82, -13, t=5.66; right: 12, -91, 8, t=4.41; 314 voxels] (Fig. 3A; Supplementary Table 3).

**Figure 3.**
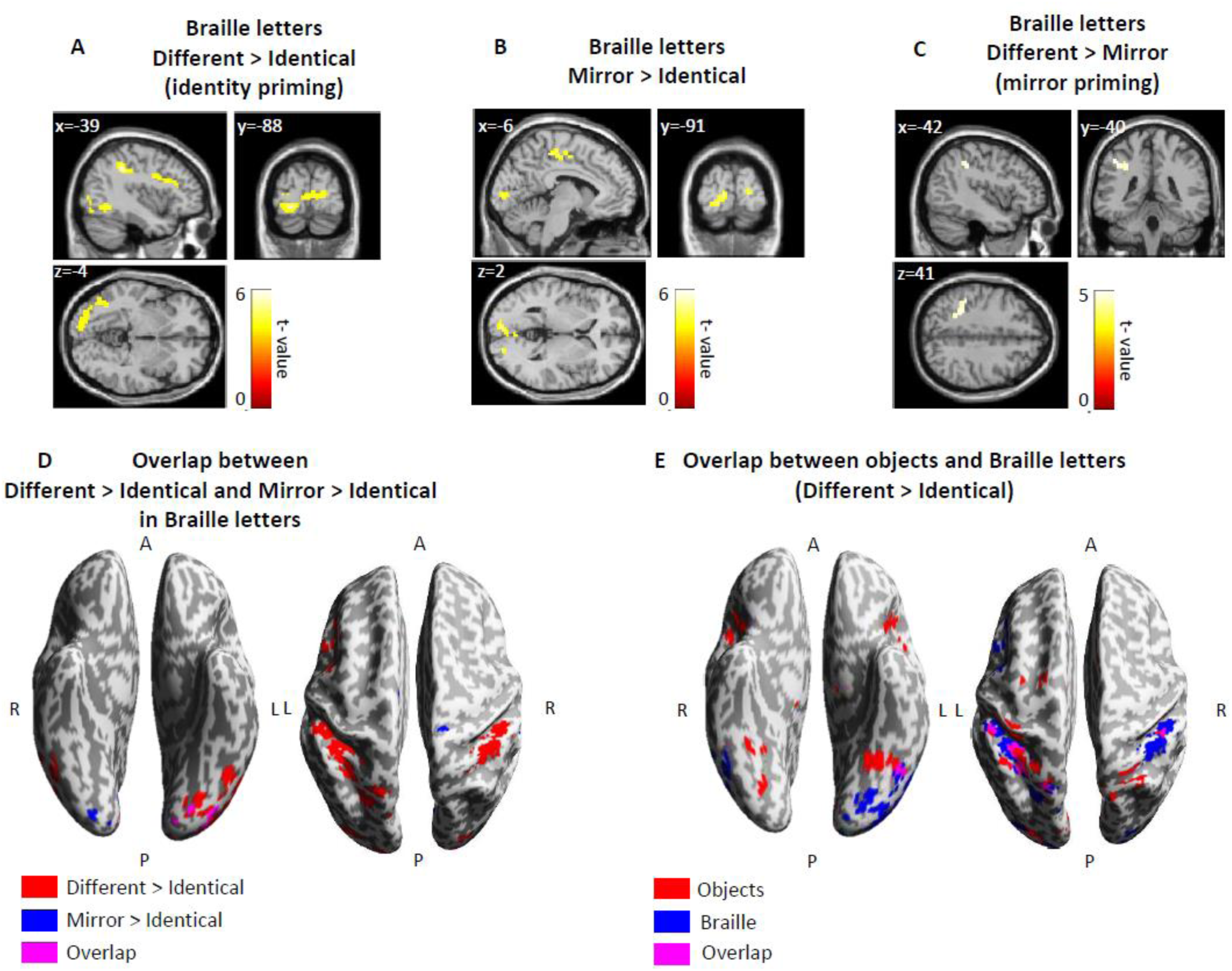
fMRI priming effects for Braille letters. **(A)** Identity priming: Braille letters induced bilateral activation in the brain’s dorsal parietal and posterior medial frontal regions. Additionally, activation was observed in the left occipito-temporal regions, particularly in the left inferior occipital gyrus, left lingual gyrus, and left middle occipital gyrus. **(B)** Mirror priming: Braille letters presented in mirror orientation induced greater activation in bilateral regions of the occipital cortex, specifically in the left calcarine and right lingual gyrus. **(C)** Absence of mirror priming: Braille letters did not produce mirror priming in occipito-temporal regions of the brain. Only the left inferior parietal lobule was activated. **(D)** Overlap between contrasts: Different > identical and mirror > identical in Braille letters revealed activation in the left inferior occipital gyrus (−21, -88, -7), illustrating how the brain distinguishes between two letters. **(E)** Activation comparison: Pairs of different Braille letters and objects evoked greater activation than pairs of the same letters and objects in the left inferior occipital gyrus (−48, -61, -13) and parietal cortex. Thresholds: **(A-D)** p < 0.001 unc. voxel-wise, p < 0.05 FWE cluster-wise; **(E)** only objects: p < 0.05 FWE voxel-wise, p < 0.05 FWE cluster-wise; R-right, L-left, A-anterior, P-posterior.

Next, we contrasted Braille letters presented in mirror > identical conditions (mirror priming), for which we expected to find similar priming effects to those observed for the different > identical pairs. We found significant activation in the right parietal cortex (postcentral gyrus: 12, -24, 62; t=5.11, 94 voxels) and bilaterally in the occipital cortex [left: inferior occipital gyrus (−21, -88, -7; t=4.55, 123 voxels); right: superior occipital gyrus (21, -88, 5; t=4.12, 49 voxels)] (Fig. 3B; Supplementary Table 3).

Finally, we contrasted Braille letters presented in different > mirror conditions, and we observed a significant activation in the left parietal lobule (−42, -40, 41; t=4.31, 57 voxels) (Fig. 3C; Supplementary Table 3).

Additionally, we found that effects for identity and mirror priming overlapped in the left inferior occipital gyrus (with the sub-peak: -21, -88, -7) (Fig. 3D). Next, we overlapped the main results of identity priming (different > identical) obtained for everyday objects and those for Braille letters, and we observed that both stimuli evoked greater activation in the left inferior occipital gyrus (with the sub-peak: -48, -61, -13) (Fig. 3E).

#### ROI analysis

First, similar to the ROI analysis for the everyday objects, we conducted repeated-measures ANOVAs (with 3 conditions: identical, mirror, and different) for Braille letters with participants’ brain activation averaged across all voxels within the anatomical masks as the dependent variable. Only in the left lateral occipital cortex we found a significant effect of conditions [F(2,34) = 10.22, p < 0.001; η2 = 0.38], with the pattern of results characteristic for breaking mirror invariance that is, we observed a significant difference between Braille letters presented in identical relative to mirror (mirror > identical p = 0.009) and different condition (different > identical p < 0.001) with no differences observed between identical and mirror condition (Fig. 4C). In the primary visual cortex (V1) and middle temporal cortex, we observed a significant effect of condition (all F’s > 3.802; all p < 0.032; η2 between 0.18 and 0.26), but it did not follow the pattern of activation where the identical condition was significantly different from both mirror and different conditions. In the remaining ROIs of the ventral visual stream, we did not observe a significant effect of orientation either in the fusiform gyrus, VWFA, primary visual cortex (V2), or dorsal extrastriate cortex (all F’s < 3.18; all p > 0.054; η2 between 0.07 and 0.16). In the parietal cortex, we found that the effect of orientation was significant only in the left parietal cortex [F(2,34) = 4.46, p = 0.019; η2 = 0.21], with significant differences observed between Braille letters presented in identical relative to mirror (mirror > identical p = 0.033) and different conditions (different > identical p < 0.042) (Fig. 4D).

## **4.** Discussion

The present study investigated whether the mirror-invariance phenomenon can be observed for tactile objects in congenitally blind subjects and whether it is specific to regions previously reported for mirror-invariant visual object recognition. Second, we aimed to determine the locus of repetition suppression for identical but not mirror Braille letters - a neural signature of breaking mirror-invariance - in congenitally blind individuals and whether it is specific to the location of the sighted VWFA.

We obtained two main findings. First, we identified brain regions that in congenitally blind individuals processed tactile objects in a mirror-invariant way. Repetition suppression for identical and mirror pairs of everyday objects was observed in parietal and occipital regions, specifically in the LOC and anterior parts of the ventral visual stream. Second, the left parietal regions and the LOC, but not the anterior vOTC (sighted VWFA), exhibited repetition suppression for identical but not mirror Braille letters - key signature of reading expertise found in the sighted.

Mirror-invariance has, so far, only been reported for visual objects (e.g., Pegado et al., 2011; Dilks et al., 2011). The repetition suppression found for identical and mirror pairs of everyday objects in regions of the occipito-temporal and parietal cortex of blind individuals — which overlaps with regions involved in mirror-invariant visual object recognition in sighted individuals — extends these results, demonstrating that this perceptual bias is not exclusive to the visual modality (Rollenhagen & Olson, 2000; Xu et al., 2023).

The sighted parieto-occipital regions are engaged in visual (Dehaene et al., 2010; Dilks et al., 2011) and tactile object recognition (Kitada et al., 2014; Fujimoto et al., 2017). In fact, parietal regions have been linked to visual and haptic object manipulation and to be particularly crucial for the control of spatially guided actions (Lucan et al., 2010; Kim & Zatorre, 2011). The latter was suggested to result from the anatomical proximity of parietal regions to sensorimotor and occipital cortices. In line with this notion, high functional connectivity between these regions has been repeatedly reported in sighted individuals (e.g., Rolls et al., 2023). In blind individuals - an even higher than in the sighted - functional connectivity of parietal and occipital regions was reported, which has been attributed to their overtrained tactile skills (Collignon et al., 2013; Qin et al., 2015; Jiao et al., 2023). Parietal and occipital areas are often simultaneously activated in blind individuals during tactile (Stilla et al., 2008), spatial imagery (Vanlierde et al., 2003), and spatial working memory (Bonino et al., 2008) tasks. The posterior parietal cortex (PPC) was suggested to integrate spatial features of an object into a multimodal high-level representation (Rolls et al., 2023), while the inferolateral occipito-temporal cortex has been found activated during shape dissemination (e.g., Xu et al., 2023), independent of input modality - if the information of the geometry is conveyed (Hannagan, 2015). In fact, Xu and colleagues (2024) have demonstrated that while parieto-occipital cortices of both blind and sighted individuals were activated during auditory processing of shape and category of manmade objects, the inferolateral occipitotemporal cortex showed higher activation in a shape verification than conceptual task (in which the participants judged the conceptual similarity of objects), corroborating the notion of Hannagan and colleagues (2015), of vOTC’s neurons being ‘apt at recognizing the shapes of objects’ (p.379) independent of the modality.

The repetition suppression for identical but not mirror Braille letters was identified in the parietal and the LOC but not in the anterior vOTC of congenitally blind individuals. Regions beyond the VWFA seem to play a significant role in visual reading. For instance, PPC was linked to grapheme-phoneme mapping, letter-identity processing (Reilhac et al., 2013), and visual working memory, especially when reading becomes more attentionally demanding, e.g., when word-forms are degraded or during spelling-like reading tasks (Deschamps et al., 2014). In blind individuals, the PPC seems the main locus of reading-related orthographic processing: Responses to Braille words in these regions were lateralized according to language processing areas but not to the reading hand (Tian et al., 2023) and the PPC’s activity increased with the number of letters in the uncontracted word version (Liu et al., 2023) suggesting its role in letter identification and letter retrieval. Our finding of repetition suppression for only identical Braille letters in the PPC corroborates these results.

The observed priming effects for letters were specific to the LOC region as well. Recently, Haut and colleagues (2024) demonstrated that while somatosensory-parietal regions and the VWFA hosted sensory and perceptual representations of tactile Braille letters, respectively, the LOC served as a “hinge” region hosting both these representations. Reconciling their results with our findings, it could be speculated that the LOC serves as the primary region for grapheme-to-phoneme mapping in tactile reading (Brem et al., 2010; Pegado et al., 2014), similar to the sighted VWFA (Pegado et al., 2014). In fact, the LOC was reported to be sensitive to linguistic properties of visual Braille and Latin words in sighted Braille readers (Cerpelloni et al., 2024) and to feature a similar activity pattern as their VWFA. Finally, an MEG study in sighted readers indicated the presence of a Letter Form Area in the LOC, where single letters are encoded prior to more complex letter combinations such as bigrams (Thesen et al., 2012; see Zhan et al., 2023 for fMRI results).

In sighted literate individuals, the neural signature of breaking mirror-invariance has been attributed to the VWFA (Dehaene et al., 2010; Pegado et al., 2011). The VWFA was suggested to emerge in regions dedicated to object recognition, as letters of many visual scripts are composed of line junctions that are apparent in the visual scene (McCandliss et al., 2003; Price & Devlin, 2011). Recent studies in sighted Braille readers, however, found selective VWFA activation in response to tactile and visual Braille alphabet, which lacks these visual characteristics (Siuda-Krzywicka et al., 2016; Cerpelloni et al., 2024) questioning the latter proposal. These findings, along with research in blind individuals, whose anterior vOTC exhibits sensitivity to the grammatical and semantic properties of auditory linguistic stimuli (e.g., Saccone, Tian, Bedny, 2024), suggested that the VWFA is primarily involved in the semantic processing of words in blind individuals as soon as semantic information is accessed (Wang et al., 2018). Our current findings might, to some extent, align with this notion: using a bottom-up task that did not require semantic information retrieval, we did not detect the neural signature of breaking mirror-invariance for Braille letters in their vOTC. In fact, our behavioral study (Korczyk et al., 2024) suggested that blind people are more efficient in filtering out task-irrelevant information and hence did not activate the semantic representation of letters to the extent that would be sufficient for the anterior vOTC activation. On the other hand, however, in our previous fMRI study (Rączy et al., 2019) using a bottom-up task as well, we observed a repetition suppression for identical but not different pairs of tactile -and not auditory-pseudowords in the anterior vOTC of blind individuals, indicating that the orthographic and not semantic processing was carried out in this region. Our current findings thus seem to suggest the LOC in blind individuals as a potential analog of the Letter Form Area identified in the sighted (see above).

Lastly, neither identity nor mirror priming effects were found in somatosensory regions. It was only observed in the different > mirror contrast, which was the only condition in which Braille letters differed in the number of dots, corroborating its role in low-level sensory processing (see Liu et al., 2023).

In the current study, we used pairs of Braille letters and everyday objects mirrored along the vertical axis. To thoroughly investigate the mirror-invariance phenomenon following congenital visual deprivation, future studies should incorporate mirror stimuli along the horizontal axis, too (De Heering et al., 2018). To evaluate the impact of object familiarity (Margalit et al., 2016), a broader range of stimuli, such as geometric figures and polynomial chains (De Heering et al., 2018), should be employed. Lastly, investigating the role of context and top-down modulation from higher-tier areas in breaking mirror-invariance for Braille letters would be of interest.

In summary, our study identified regions involved in the mirror-invariant processing of haptic objects in blind individuals. Among these regions, the parietal and lateral occipital cortices also exhibited neural patterns indicative of breaking mirror invariance for Braille letters. While these regions are part of a co-activated network engaged in tactile object recognition in blind individuals, we hypothesize they serve distinct functions. The parietal regions likely process letter features such as orientation, serving as an initial step in distinguishing different and mirror letters as distinct. In contrast, the LOC integrates shape information with letter identity representation, possibly serving as an early stage in grapheme-to-phoneme conversion. The neural adaptation for identical but not mirror pairs of Braille letters was notably absent in the anterior vOTC. This suggests that its role in reading in blind individuals might be notably different from the one in the sighted.

## **5.** Author Contributions

Author contributions: M.K., K.R., and M.S. designed research; M.K. performed research; M.K. analyzed data; M.K., K.R., and M.S. wrote the paper.

## 6. Data Availability Statement

Data will be made available on an open-access server upon publication on the research article and are available upon request to the reviewers.

## 7. Conflict of interest statement

The authors declare that the research was conducted in the absence of any commercial or financial relationships that could be construed as a potential conflict of interest.

### Acknowledgments

This work was supported by the Polish National Science Centre (Grant Number 2018/30/A/HS6/00595) to M.S. We would like to thank all of our participants for their efforts and for participating in the study.

