## Supplementary for "Repetition suppression for mirror images of objects and not Braille letters in the ventral visual stream of congenitally blind individuals"

**Table 1.** Brain activation induced by everyday objects in contrast: different >identical

| Anatomical region | BA | Cluster size (voxels) | MNI-coordinates X | Y | Z | t-value |
|---|---|---|---|---|---|---|
| different > identical everyday objects | | | | | | |
| L fusiform gyrus | 37 | 75 | -33 | -55 | -10 | 7.34 |
| L fusiform gyrus | 37 | | -33 | -55 | -19 | 6.47 |
| L inferior occipital gyrus | 37 | | -48 | -61 | -13 | 5.33 |
| R insula lobe | 13 | 114 | 33 | 20 | 5 | 6.98 |
| R insula lobe | 13 | | 39 | 17 | -4 | 6.13 |
| L inferior parietal lobule | 40 | 367 | -39 | -40 | 50 | 6.98 |
| L inferior parietal lobule | 7 | | -33 | -52 | 50 | 6.71 |
| L inferior parietal lobule | 40 | | -48 | -28 | 44 | 6.46 |
| R insula lobe | 13 | 97 | -36 | 17 | 5 | 6.62 |
| R posterior-medial frontal | 6 | 135 | 6 | 11 | 53 | 6.58 |
| L posterior-medial frontal | NA | | 0 | 2 | 56 | 5.79 |
| R supplamentary motor cortex | 8 | | 6 | 20 | 41 | 5.20 |
| thalamus | NA | 27 | -9 | -19 | -4 | 6.37 |
| thalamus | NA | | -3 | -22 | -10 | 5.67 |
| R precuneus | 7 | 45 | 12 | -70 | 44 | 6.26 |
| thalamus | NA | 33 | 9 | -16 | -4 | 6.25 |
| thalamus | NA | | 6 | -28 | -7 | 5.25 |
| L middle occipital gyrus | 18 | 34 | -24 | -88 | 8 | 6.11 |
| R caudate nucleus | NA | 17 | 9 | 8 | 5 | 6.09 |
| L cuneus | 18 | 57 | -3 | -76 | 20 | 5.95 |
| L lingual gyrus | 17 | | 0 | -82 | 5 | 5.81 |
| L cuneus | 19 | | -6 | -85 | 26 | 5.36 |
| R superior parietal lobule | NA | 60 | 24 | -52 | 56 | 5.85 |
| R superior parietal lobule | 7 | | 24 | -61 | 59 | 5.84 |
| R superior parietal lobule | 7 | | 30 | -61 | 53 | 5.51 |
| L superior frontal gyrus | 6 | 31 | -24 | -7 | 53 | 5.64 |
| R postcentral gyrus | 1 | 21 | 48 | -31 | 53 | 5.48 |
| R supramarginal gyrus | 1 | | 45 | -31 | 44 | 5.18 |
| R fusiform gyrus | 19 | 14 | 27 | -70 | -7 | 5.44 |
| R fusiform gyrus | 37 | 12 | 30 | -52 | -13 | 5.09 |

NA. not appicable. Thresholds: $p < 0.05$ FWE voxel-wise. $p < 0.05$ FWE cluster-wise.

**Table 2.** Brain activation induced by everyday objects in contrast: different > mirror

| Anatomical region | BA | Cluster size (voxels) | MNI-coordinates | | | t-value |
|---|---|---|---|---|---|---|
| | | | X | Y | Z | |
| different> mirror everyday objects | | | | | | |
| L postcentral gyrus | 1 | 586 | -45 | -28 | 50 | 7.47 |
| R precuneus | 7 | | 12 | -70 | 47 | 7.31 |
| L inferior parietal lobule | 40 | | -39 | -37 | 50 | 7.03 |
| R posterior-medial frontal | 6 | 257 | 6 | 5 | 53 | 6.95 |
| L posterior-medial frontal | 6 | | -3 | -7 | 56 | 6.82 |
| L middle cingulate gyrus | 24 | | -3 | 2 | 41 | 5.88 |
| brain stem | NA | 146 | 6 | -28 | -7 | 6.63 |
| Thalamus | NA | | 12 | -16 | -4 | 6.09 |
| Thalamus | NA | | 6 | -16 | 5 | 5.51 |
| Thalamus | NA | 155 | -12 | -28 | -4 | 6.46 |
| Thalamus | NA | | -9 | -22 | 2 | 6.36 |
| Thalamus | NA | | -15 | -19 | 8 | 6.29 |
| L fusiform gyrus | 37 | 201 | -33 | -55 | -13 | 6.45 |
| L fusiform gyrus | 18 | | -24 | -73 | -13 | 5.98 |
| L cerebelum (VI) | NA | | -30 | -70 | -19 | 5.68 |
| L insula lobe | 13 | 62 | -39 | 14 | -1 | 6.41 |
| L insula lobe | 45 | | -30 | 20 | 8 | 6.26 |
| L insula lobe | 13 | | -45 | 5 | 2 | 5.08 |
| R postcentral gyrus | 1 | 134 | 48 | -28 | 53 | 6.38 |
| R postcentral gyrus | 1 | | 51 | -22 | 44 | 5.84 |
| R postcentral gyrus | 1 | | 33 | -31 | 50 | 5.51 |
| L rolandic operculum | 40 | 86 | -48 | -22 | 17 | 6.37 |
| L rolandic operculum | 13 | | -33 | -28 | 17 | 5.41 |
| R insula lobe | 13 | 74 | 33 | 17 | 5 | 6.30 |
| R insula lobe | 13 | | 42 | 14 | -1 | 5.62 |
| R cerebelum (IV-V) | NA | 115 | 12 | -55 | -16 | 5.87 |
| R Cerebelum (VI) | NA | | 27 | -49 | -25 | 5.58 |
| cerebellar vermis (6) | NA | | 0 | -70 | -16 | 5.34 |
| L precentral gyrus | 6 | 13 | -21 | -13 | 68 | 5.71 |
| L superior frontal gyrus | 6 | | -24 | -7 | 53 | 5.24 |
| R caudate nucleus | NA | 18 | 12 | 8 | 8 | 5.59 |
| R putamen | NA | 15 | 33 | -22 | -1 | 5.52 |
| L cuneus | 18 | 17 | -3 | -76 | 23 | 5.28 |
| L cuneus | 19 | | -6 | -82 | 32 | 4.89 |

NA. not appicable. Thresholds: $p < 0.05$ FWE voxel-wise. $p < 0.05$ FWE cluster-wise.

**Table 3.** Brain activation induced by Braille letters in three contrasts: different>identical, mirror>identical, different>mirror.

| Anatomical region | BA | Cluster size (voxels) | MNI-coordinates | | | t-value |
|---|---|---|---|---|---|---|
| | | | X | Y | Z | |
| different>identical Braille letters | | | | | | |
| L inferior parietal lobule | 40 | 372 | -42 | -40 | 41 | 6.00 |
| L inferior parietal lobule | NA | | -33 | -43 | 41 | 5.65 |
| L superior parietal lobule | NA | | -21 | -61 | 50 | 5.34 |
| L lingual gyrus | 18 | 314 | -30 | -82 | -13 | 5.66 |
| L inferior occipital gyrus | 18 | | -21 | -88 | -7 | 5.08 |
| R cuneus | 18 | | 21 | -94 | 8 | 4.51 |
| R inferior parietal lobule | 40 | 158 | 45 | -37 | 47 | 5.49 |
| R inferior parietal lobule | 40 | | 42 | -31 | 35 | 4.41 |
| R supperior parietal lobule | 40 | | 33 | -40 | 41 | 4.40 |
| L inferior frontal gyrus | 44 | 128 | -51 | 14 | 29 | 4.88 |
| L inferior frontal gyrus | 6 | | -42 | 5 | 29 | 4.36 |
| L inferior frontal gyurs | 44 | | -42 | 17 | 26 | 3.95 |
| L inferior temporal gyrus | 19 | 70 | -42 | -64 | -7 | 4.53 |
| L middle occipital gyrus | 19 | | -45 | -70 | -1 | 4.13 |
| L posterior-medial frontal | 6 | 51 | -6 | 11 | 50 | 4.46 |
| R posterior-medial frontal | 6 | | 9 | 14 | 50 | 4.14 |
| L superior medial gyrus | 8 | | -3 | 23 | 41 | 3.63 |
| mirror>identical Braille letters | | | | | | |
| R postcentral gyrus | NA | 94 | 12 | -34 | 62 | 5.11 |
| R posterior-medial frontal | 6 | | 6 | -25 | 56 | 4.52 |
| L paracentral lobule | 6 | | -3 | -25 | 53 | 4.19 |
| L inferior occipital gyrus | 18 | 123 | -21 | -88 | -7 | 4.55 |
| L calcarine gyrus | 17 | | -9 | -91 | 2 | 4.11 |
| L calcarine gyrus | 17 | | -3 | -85 | 2 | 3.93 |
| R superior occipital gyrus | NA | 49 | 21 | -88 | 5 | 4.12 |
| R lingual gyrus | 18 | | 21 | -85 | -4 | 3.97 |
| R lingual gyrus | 18 | | 30 | -79 | -1 | 3.60 |
| different> mirror Braille letters | | | | | | |
| L inferior parietal lobule | 40 | 57 | -42 | -40 | 41 | 4.31 |
| L inferior parietal lobule | BA | | -30 | -43 | 41 | 4.18 |
| L inferior parietal lobule | 7 | | -27 | -52 | 44 | 3.80 |

NA. not appicable. Thresholds: $p < 0.001$ unc. voxel-wise. $p < 0.05$ FWE cluster-wise

**A The conveyor belt**

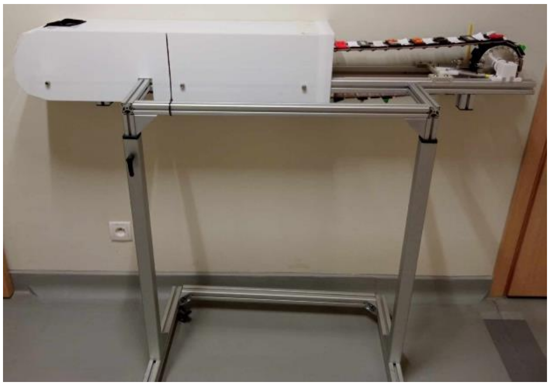

**B Plastic plates and spacers**

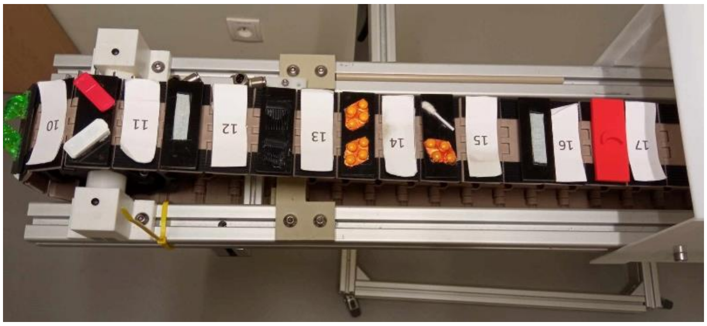

**C Location of the conveyor belt inside the scanner**

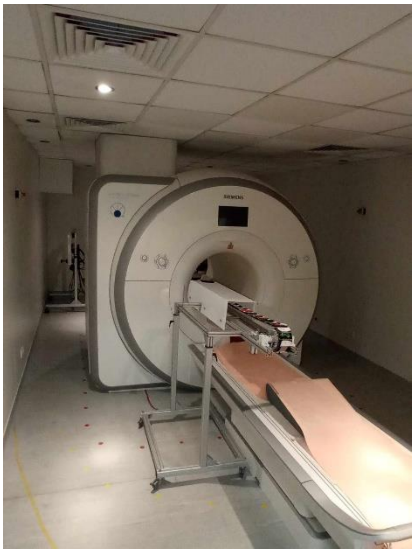

**D aperture with the pairs of stimuli**

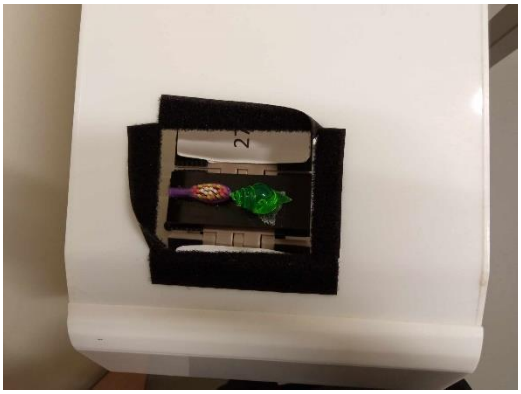

**Supplementary Figure 1. (A)** The conveyor belt was specially designed for this study to be fMRI-compatible. **(B)** A picture of the belt with plastic plates containing pairs of stimuli and spacers – gaps between each stimuli plate intended to avoid any possible mistakes. **(C)** The device was placed above the participants' thighs on their reading side. The researcher moved the chain and remained at the end of the device. **(D)**The participants' reading hand was located in a special aperture, allowing them to touch only one pair of stimuli at a time.

**A Anatomical masks for the left occipital-temporal** **B Anatomical masks for the bilateral parietal cortex**

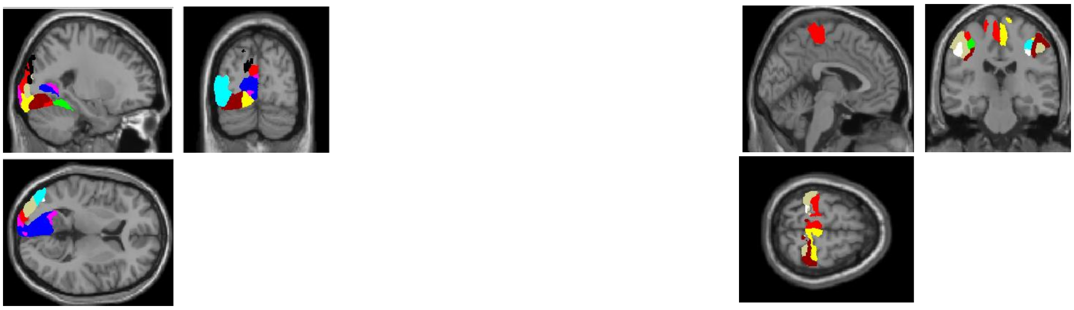

**Supplementary Figure 2. (A)** Anatomical masks for the left occipital-temporal were created using the SPM Anatomy Toolbox 2.2b (Eickhoff et al., 2005). The left occipital-temporal regions encompassed areas such as 1) the primary visual cortex (V1 and V2 i.e., BA 17 and BA 18), 2) middle-temporal cortex (hOC5 (V5 / MT+))(, 3) ventral (V3v / V4) and 4) dorsal extrastriate cortex (hOC3d / hOC4d), 5) fusiform gyrus (Areas FG1, FG2, FG3 and FG4), and 6) lateral occipital cortex (extrastriate areas hOc4la and hOc4lp). **(B)** Anatomical masks for the bilateral parietal cortex created using the SPM Anatomy Toolbox 2.2b (Eickhoff et al., 2005) included: 1) the intraparietal sulcus(Areas hIP1, hIP2, hIP3), 2) motor cortex (Areas 4a and 4p), and 3) primary somatosensory cortex (Areas 1, 2, 3a, 3b).
